# First-trimester exposure to anti-cytomegalovirus medications: a study of the effects of valaciclovir, letermovir and maribavir on early placental development

**DOI:** 10.1101/2025.05.01.651765

**Authors:** Ishara Atukorala, Natasha de Alwis, Sally Beard, Swetha Raghavan, Natalie J. Hannan, Lisa Hui

## Abstract

Congenital cytomegalovirus (CMV) infection is the leading preventable cause of childhood disability, including sensorineural hearing loss and cerebral palsy in high-income countries. Early maternal treatment with valaciclovir has been shown to reduce fetal infection. However, newer, more potent anti-CMV medications such as letermovir and maribavir are not routinely used in pregnancy due to limited safety data. This study aimed to assess the effect of aciclovir (the active metabolite of valaciclovir), letermovir, and maribavir on cell viability, functionality, and key cell signalling pathways in first-trimester human placental tissue.

First-trimester placental tissue was obtained from individuals undergoing surgical abortion for psychosocial indications at 9-13 weeks of gestation. The effect of anti-CMV medications on primary cytotrophoblast cell survival was measured using a standard cell viability assay. The effects of anti-CMV medication exposure on Wnt and epithelial-to-mesenchymal transition (EMT) signalling pathways were evaluated in placental explant cultures. Their impact on cellular proliferation and migration was assessed using placental outgrowth cultures. Exposure to physiological or supraphysiological levels of aciclovir, letermovir, or maribavir for 48 hours did not significantly alter cytotrophoblast cell viability compared to vehicle-treated controls. Aciclovir did not significantly impact Wnt and EMT signalling gene expression in first-trimester placental explants; however, letermovir and maribavir induced significant gene dysregulation. None of the three anti-CMV medications altered first-trimester trophoblast proliferation or migration.

These results provide reassuring data supporting the safety of maternal valaciclovir therapy in the first trimester. They may also inform the selection of newer anti-CMV medications for future clinical trials for use in pregnancy.

## Introduction

Cytomegalovirus (CMV) is the most common cause of congenital infection with an estimated birth prevalence of 0.5% in high-income countries (1). Maternal infection during pregnancy can lead to fetal transmission, resulting in severe outcomes such as stillbirth, neonatal death, and long-term neurological disabilities, including sensorineural hearing loss, cerebral palsy, and neurodevelopmental delay (2). High-dose valaciclovir treatment has been shown to reduce the risk of fetal infection by 70% when administered after maternal primary CMV infection in early pregnancy (3-6). International health authorities now recommend an oral valacyclovir dosage of 8 g/day for maternal primary infection during the periconceptional period or in the first trimester of pregnancy (7). However, newer and more potent antiviral agents, such as letermovir and maribavir, are not currently recommended in pregnancy due to limited safety and efficacy data (8).

The placenta plays a central role in fetal development by facilitating nutrient exchange, waste removal, hormone production, and immune modulation (9, 10). The first trimester is a critical period for placental and fetal development, during which disruptions may have long-term consequences for pregnancy outcomes. Assessing the potential effects of anti-CMV therapies on placental development is essential for determining their safety when used in early pregnancy to prevent fetal infection.

Early placental development is driven by several key cell signalling pathways (9, 11). The Wnt signalling pathway plays a crucial role in the regulation of cell proliferation, migration, and invasion (12-14). Wnt signalling is essential for the transition of trophoblast cells from a proliferative to an invasive phenotype (15). Dysregulation of this pathway has been implicated in pregnancy complications such as miscarriage (13), congenital abnormalities and pre-eclampsia (16, 17).

Another critical process in placental development is epithelial-to-mesenchymal transition (EMT) (18, 19). Through EMT, differentiating extravillous cytotrophoblasts gain migratory and invasive properties, facilitating their invasion into the maternal decidua basalis to remodel the maternal uterine spiral arteries (20, 21). This process is essential for establishing adequate placental perfusion (22).

Despite the growing acceptance of valaciclovir as a therapy to reduce fetal infection (7, 23), there is a paucity of data on its effects on the first-trimester human placenta. Current knowledge is based on data derived from third-trimester human placentas or transformed trophoblast cell lines that may not be applicable *in vivo* (24, 25).

This study aimed to assess the effects of aciclovir (the active metabolite of valaciclovir), letermovir, and maribavir on first-trimester placental development. Specifically, we examined their impact on cytotrophoblast viability, the expression of Wnt and EMT signalling markers, and placental villous outgrowth.

## Methods

### First-trimester placenta sample collection

Ethical approval for placental sample collection was obtained from the Austin Health Human Research Ethics Committee (Ref. No. 68965). Patients provided written informed consent prior to the collection of placental tissue samples.

Placental tissue was obtained according to previously established protocols (26). In brief, the tissue was collected from individuals undergoing elective first-trimester surgical termination of live singleton pregnancies under general anaesthesia via curettage, or a combination of aspiration and curettage. Abortions were performed for maternal psychosocial indications; pregnancies with complications including suspected CMV infection or aneuploidy were excluded. Placental tissue was separated from other conceptus material and then washed in phosphate-buffered saline (PBS; 137 mM NaCl, 10 mM Na_2_HPO_4_, 1.8 mM KH_2_PO_4_, 2.7 mM KCl, pH 7.4). Placental tissue was stored in cold Dulbecco’s Modified Eagle Medium (DMEM) Glutamax (Gibco^TM^; ThermoFisher Scientific, Scoresby, VIC, Australia) for a short period (0.5-3 hours) before dissection for subsequent assessment.

### Cytotrophoblast isolation and culture

Primary human cytotrophoblast cells were isolated using an established protocol (27), with the following modifications for first-trimester tissue: whole placental tissue sample was placed in a petri dish and placental tissue gently dissected away from the fetal membranes (yielding 2-9 g total dissociated tissue), and placed in DMEM Glutamax for three, 10-minute incubations with 0.25% trypsin (ThermoFisher Scientific) and 1% DNase (Sigma Aldrich, St Louis, MO, USA) at 37°C to digest tissue. Percoll centrifugal separation of the cells and CD9 negative selection was performed as previously described (27).

### Cytotrophoblast cell viability assay

Primary cytotrophoblast cells were plated at 25,000 cells/well in DMEM Glutamax supplemented with 10% fetal bovine serum (FBS; ThermoFisher Scientific) and 1% Antibiotic-Antimycotic (AA; ThermoFisher Scientific) on fibronectin-coated 96-well cell culture plates (n=5, 2 technical replicates per treatment) and incubated at 37°C, 8% O_2_, 5% CO_2_ overnight. The cells were then treated with aciclovir (1, 10, 25, 50 and 100 µM; Tokyo Chemical Industries, Tokyo, Japan), letermovir (0.1, 1, 10, 25 and 50 µM; Sigma Aldrich), maribavir (0.1, 1, 10, 25 and 50 µM; Sigma Aldrich), or dimethyl sulfoxide (1% v/v; Sigma Aldrich) as the vehicle control. Aciclovir is the metabolite of valaciclovir. After internalisation, the small intestine and liver convert valaciclovir to aciclovir, which then enters the bloodstream (28). Since this conversion is absent in our experimental models, we have used the metabolised product aciclovir.

Cells were incubated with the treatments for 48 hours at 37°C, 8% O_2_, 5% CO_2_. Cell viability was assessed using the CellTiter 96 AQueous Non-Radioactive Cell Proliferation Assay (MTS) assay according to the manufacturer’s instructions (Promega, Madison, WI, USA) and optical density was analysed using a FluoStar Omega^TM^ Microplate Reader. The statistical analysis was performed with a mixed-effects analysis with Dunnett’s correction, using GraphPad Prism ^TM^ software (version 10.1.0 (316).

### Placental explant culture

Placental tissue explants (∼30 mg) were dissected from first-trimester placental tissue and cultured in DMEM/10% FBS/1% Antibiotic-Antimycotic media at 37°C, 8% O_2_, 5% CO_2_ for 24 hours before treatment. After replacement with fresh media, explants (n=6, 3 technical replicates per treatment) were treated with aciclovir (10-100 µM), letermovir (1-50 µM), maribavir (1-50 µM) or DMSO (1% v/v) as the vehicle control and incubated for a further 48 hours before collecting the tissue for RNA or protein extraction.

### Quantitative RT-PCR for WNT and EMT signalling marker genes

RNA was extracted from explants using Sigma GenElute™ Total RNA Purification Kit, following the manufacturer’s protocol. RNA was quantified using LVis Plate in FluoStar Omega Microplate Reader. RNA (0.2 µg) was used for cDNA synthesis using High-Capacity cDNA reverse transcription kit (Applied Biosystems™) according to the manufacturer’s protocol, in a MiniAmp Thermal Cycler (ThermoFisher Scientific). The gene expression of Wnt signalling markers MYC proto-oncogene (MYC) and cyclin D1 (CCND1), EMT signalling markers Vimentin (VIM) and Snail family transcriptional repressor 1 (SNAI1), and reference genes DNA topoisomerase 1 (TOP1) and Cytochrome C1 (CYC1), was analysed using TaqMan Gene Expression Assay. RT-PCR was performed using TaqMan Fast Advanced Master Mix (Applied Biosystems™) and QuantStudio^TM^ 5 Real-Time PCR System with the following run conditions: 50 °C for 2 min; 95 °C for 2 min, 95 °C for 1 s, 60 °C for 20 sec (40 cycles). All data were normalized to the reference genes *TOP1* and *CYC1* as internal controls and calibrated against the average Ct of the control samples. All cDNA samples were run in duplicate. Nested one-way ANOVA was used for statistical analysis (GraphPad Prism software).

### Western blotting for Wnt and EMT signalling markers

Explant tissue was homogenised in RIPA lysis buffer (150 mM NaCl, 50 mM Tris pH 8.0, 0.1% (w/v) SDS, 1% (v/v) Triton X 100, 0.5% (v/v) sodium deoxycholate) with freshly added protease inhibitors and phosphatase inhibitors, on ice. The homogenate was centrifuged at 10,000 g for 20 min to remove cell debris. Equal protein amounts (30 µg) were heated with 4x Laemmli buffer (8% (w/v) SDS, 10% (v/v) glycerol, 200mM Tris-HCL pH 6.8 and trace of bromophenol blue), and 2M DTT, at 95°C for 2 min and separated on gradient 4-15% polyacrylamide gels (BioRad). Gels were electrophoresed at 120 V for about 90 min. Proteins were transferred to PVDF membranes (ThermoFisher Scientific) and blocked with 5% skim milk. Membranes were incubated with primary antibodies for β-catenin (CTNNB1; Cell signalling) as a Wnt signalling marker, E-cadherin (Cell signalling) as an EMT marker or β-actin (Cell signalling) as the loading control, at 4°C overnight. After washing the access antibody off, the membranes were incubated with the relevant secondary antibodies for 1 h at room temperature. Membranes were imaged using ChemiDoc (Bio-Rad). Densitometric analysis of the Western blots was performed using Image Lab Software (Bio-Rad). E-cadherin and β-catenin levels in each sample were normalised to the β-actin expression of the same sample. Normality of the data distribution was determined using the Shapiro-Wilk test. Each treatment was compared to the DMSO control using the Student’s t-test or the Mann-Whitney U-test to determine statistical significance. All tests were conducted in GraphPad Prism 10.1.0.

### Placental outgrowth culture

Villous tips (3 -4mm) obtained from first trimester placental tissue (n=4) were dissected and placed into cold 24-well cell culture plates coated with 1mg/ml Cultrex Rat Collagen I (R&D Systems) in DMEM Glutamax (buffered with 5M NaOH). Collagen was allowed to set at room temperature for 15 minutes, followed by further incubation at 37°C under 5% O_2_ and 5% CO_2_ for 1 hour before addition of villous tips (1 per well). Fresh media (DMEM Glutamax, 10% FBS, 1% Antibiotic-Antimycotic) was added to each well and incubated at 5% O_2_ and 5% CO_2_, and 37°C. Following overnight incubation, images of the explants were taken using a light microscope. Media in each well was replaced with fresh media containing aciclovir (50 µM), letermovir (25 µM), maribavir (25 µM) or DMSO (1% v/v) as the vehicle control. Plates were incubated for 72 h before final images were captured to assess treatment effects.

### Image analysis and measurement of placental villous outgrowth

Each image was converted to an 8-bit format to optimise visualisation. For area measurement, the freehand selections tool was used to trace the outgrowth area manually. Once the area was outlined, the measurement was performed by selecting the analyse measurement function. The FIJI ImageJ Grid/Collection Stitching plugin was used to stitch multiple images at the final timepoint, as previously described (29). The same procedure as outlined above for individual images was followed to measure the area of the fused image. Student’s T-test was used for statistical analysis.

## Results

### Aciclovir, letermovir and maribavir treatments did not alter primary cytotrophoblast cell viability

We first used an *in-vitro* model of isolated pure primary cytotrophoblast cells to assess drug cytotoxicity, employing an MTS assay to measure cellular metabolic activity, as a measure of cell viability. There were no significant changes in the metabolic activity/viability of cytotrophoblast cells following 48-hour exposure to all concentrations of the anti-CMV medications compared to control (Figure 1).

**Figure 1.**
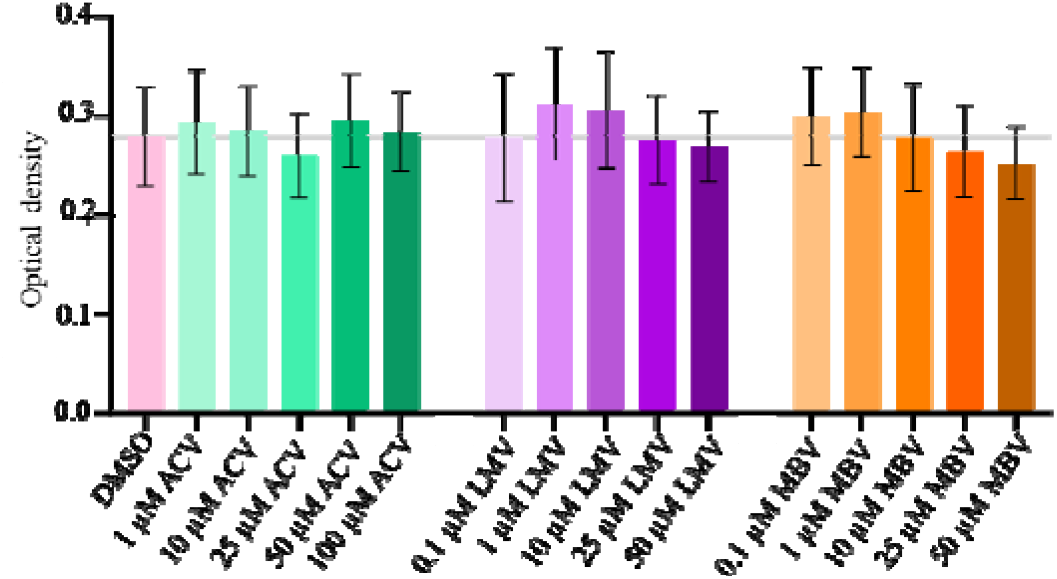
Effect of anti-CMV medications on cell viability in primary cytotrophoblasts. Primary cytotrophoblasts were isolated and plated at 25,000 cells/well. Optical density was recorded as a measure of cell viability following a 48-hour treatment with aciclovir (1, 10, 25, 50 and 100 µM), letermovir (0.1, 1, 10, 25 and 50 µM) or maribavir (0.1, 1, 10, 25 and 50 µM). DMSO (1%) was used as the vehicle control. Mean ± SEM. *n*=5 ACV: aciclovir, LMV: letermovir, MBV: maribavir

### Letermovir and maribavir but not Aciclovir impacted the Wnt signaling pathway in first-trimester placental tissue

Next, we analysed Wnt signalling activity in the presence of anti-CMV medications using a placental explant *ex-vivo* model. Aciclovir treatment did not alter placental mRNA expression of cyclin-D1 (Fig 2A) and C-MYC (n=6) (Fig 2B), or protein levels of β-catenin (n=3) (Fig 2C and 2D).

**Figure 2.**
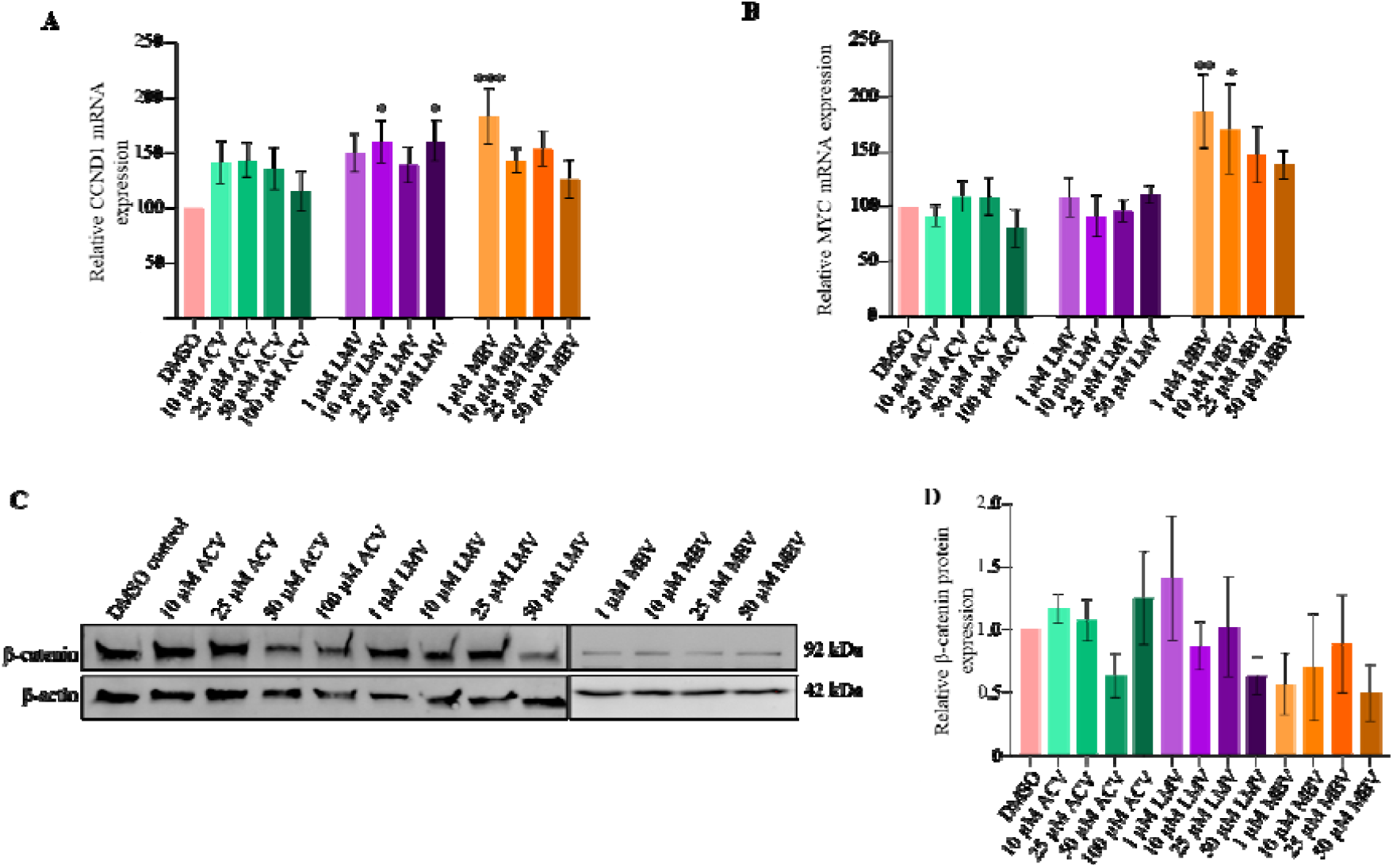
The effect of anti-CMV medications on the Wnt signalling pathway in the first-trimester placenta. Placental explants (30 mg) were treated with aciclovir (10, 25, 50, and 100 µM), letermovir (1, 10, 25, and 50 µM), maribavir (1, 10, 25, and 50 µM), or vehicle control. Transcript levels of (A) *CCND1* and (B) *MYC* were quantified using qPCR and normalised to the reference gene *TOP1*, as shown in the graphs (*n*=6). All qPCR data is presented as relative expression. (C) β-catenin protein levels are illustrated by the Western blot (representative image, *n*=3). (D) Densitometric analysis of β-catenin bands, normalised to β-actin, is presented as percentage expression relative to the DMSO control. Mean ± SEM. *p < 0.05 **p < 0.01, ***p < 0.001 ACV: aciclovir, LMV: letermovir, MBV: maribavir

Exposure to letermovir at concentrations of 10 and 50 µM significantly altered placental *CCND1* gene expression (Figures 2A) but not *MYC* or β-catenin (Fig 2C). Other letermovir concentrations did not affect *CCND1* and *MYC* gene expression or cellular β-catenin protein levels.

Maribavir treatment at 1 µM significantly increased mRNA expression of *CCND1* but not at other concentrations (Fig 2A). Maribavir at concentrations 1 and 10 µM led to a significant increase in the RNA levels of the *MYC* proto-oncogene (Fig 2B). However, none of the Maribavir treatments significantly altered the placental β-catenin protein levels.

### Aciclovir and letermovir did not alter EMT signalling in first-trimester placental tissue

We studied EMT markers, *VIM* (Fig 3A), *SNAI1* (n=6) (Fig 3B), and E-cadherin (CDH1) (n=3) (Fig 3C and 3D) using RT-PCR and Western blotting. Overall, aciclovir and letermovir did not impact the markers of EMT signalling pathway. mRNA profile of *VIM* remained unchanged with all treatment concentrations of maribavir. However, *SNAI1* mRNA expression was significantly increased after treatment with 1 µM maribavir.

**Figure 3.**
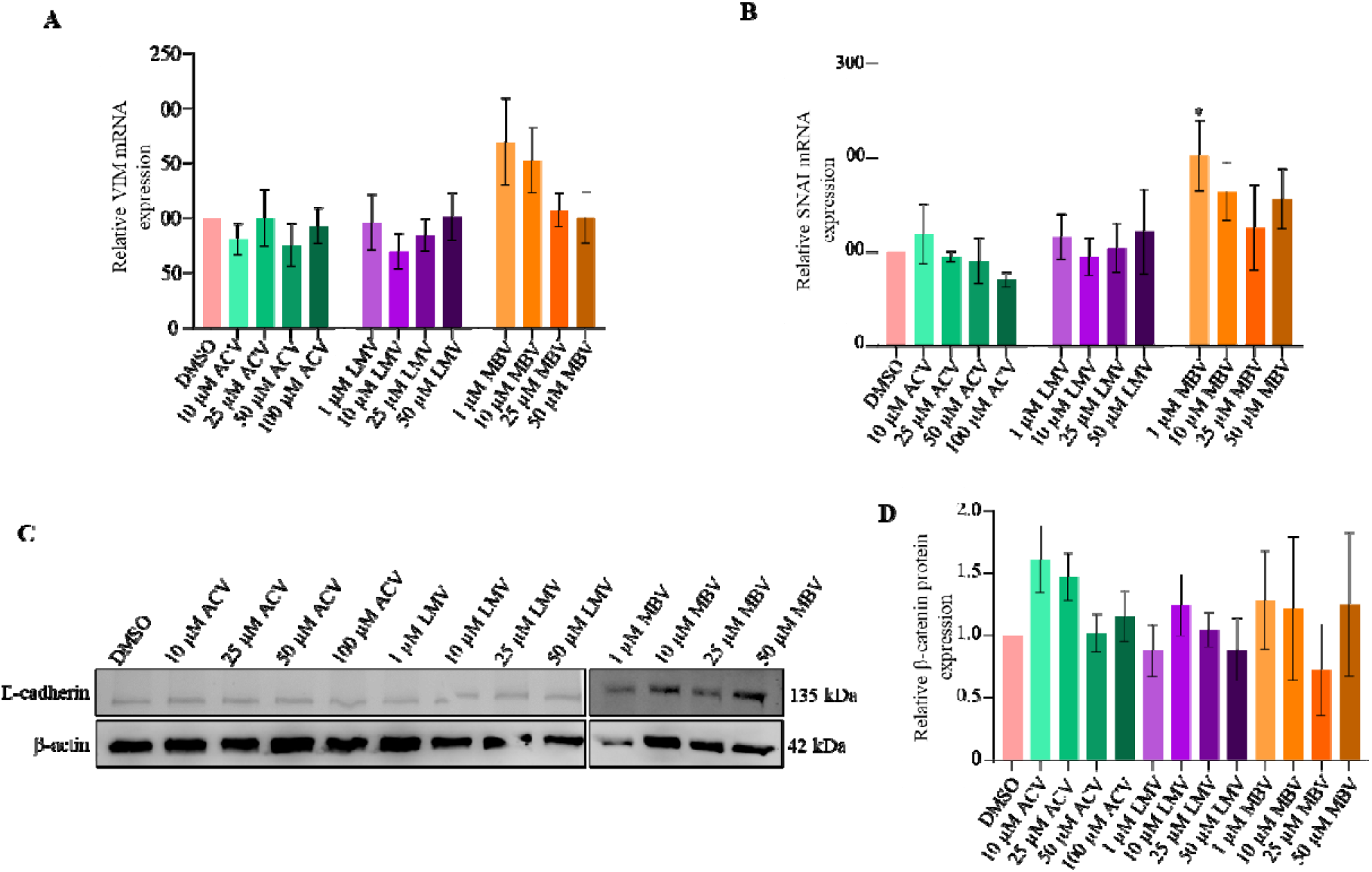
The effect of anti-CMV medications on the EMT signalling pathway in the first-trimester placenta. Placental explants (30 mg) were treated with aciclovir (10, 25, 50, and 100 µM), letermovir (1, 10, 25, and 50 µM), maribavir (1, 10, 25, and 50 µM), or vehicle control. Transcript levels of (A) Vimentin and (B) Snail were quantified using qPCR and normalised to the housekeeping gene *TOP1*, as shown in the graphs (*n*=6). All qPCR data is presented as relative expression. (C) The Western blot illustrates E-cadherin protein levels (representative image, *n*=3). (D) Densitometric analysis of E-cadherin bands, normalised to β-actin, is presented as percentage expression relative to the DMSO control, mean ± SEM. *n*=6 *p < 0.05 ACV: aciclovir, LMV: letermovir, MBV: maribavir

### Aciclovir, letermovir and maribavir do not affect first-trimester placental outgrowth

We used a placental outgrowth model to assess the impact of the antiviral treatments on placental outgrowth on collagen, as an indicator of cell proliferation and migration. We used the same physiologically relevant concentrations of aciclovir, letermovir and maribavir used in the previous experiments.

After 72 hours of treatment, the placental cells proliferated and migrated into the collagen matrix, expanding from the villous tips. Tissue outgrowths in Figure 4A are representative of the different treatment conditions. We did not observe significant changes in the outgrowth area of the placental tissue following the antiviral treatments compared to the vehicle control (Figure 4B).

**Figure 4.**
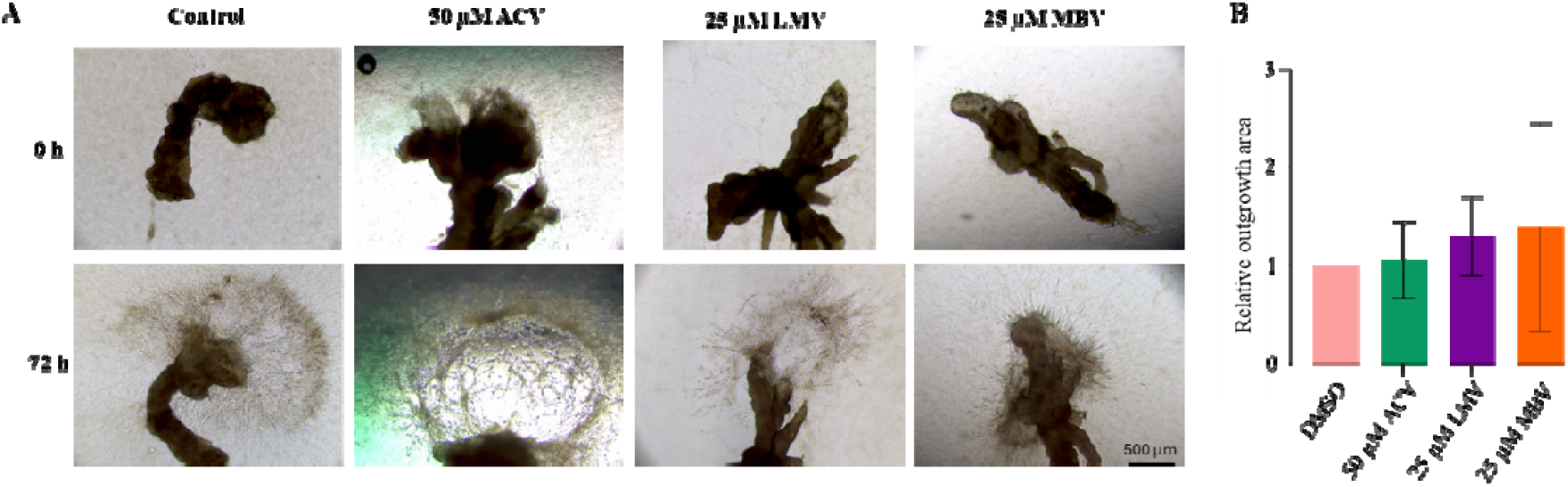
Placental villi outgrowth in response to anti-viral treatments. (A) Representative microscopy images of placental villi tips incubated in collagen-coated plates with anti-viral treatments or vehicle control for 72 hours. Following treatment, placental cells proliferated and invaded the collagen matrix, extending from the villi tips. Images were analysed using FIJI ImageJ software. (B) Quantification of the outgrowth area, measured using FIJI ImageJ, is represented as a relative value compared to the control. mean ± SEM. *n*=4 ACV: aciclovir, LMV: letermovir, MBV: maribavir

## Discussion

Our study provides the first data on the impact of three anti-CMV medications—aciclovir (the active metabolite of valaciclovir), letermovir, and maribavir—on first-trimester placental development using primary human tissues. We found that exposure to aciclovir, letermovir, or maribavir did not compromise primary cytotrophoblast viability or affect placental proliferation and migration. Aciclovir had no significant impact on Wnt or EMT signalling pathways, both of which play essential roles in placental development. However, letermovir and maribavir did induce dysregulation of selected genes in these signalling pathways, warranting further investigation. However, despite these impacts on gene expression, neither letermovir nor maribavir impaired placental viability or outgrowth *in vitro*.

Our findings are consistent with earlier studies reporting on the safety of acyclovir, letermovir and maribavir in term placental explants and trophoblast cell lines (25). However, these models may not accurately reflect the physiology of the first-trimester placenta, which represents the most clinically relevant window for the prevention of fetal infection. By contrast, our study specifically investigated first-trimester placental tissue, an especially critical period for fetal organogenesis and placental development. Our results, therefore, fill a critical knowledge gap by providing the first data on the effects of anti-CMV medications in first-trimester primary human placenta. Given that early placental development is particularly sensitive to disruptions in Wnt and EMT pathways, our findings contribute important preclinical evidence supporting the relative safety of valaciclovir in the most clinically relevant window of human development while highlighting potential areas of concern for next-generation antiviral therapies.

Valaciclovir and its metabolite, aciclovir, have been widely used in the third trimester for the suppression of genital herpes. However, the use of valaciclovir for preventing fetal CMV infection after maternal primary infection requires a substantially higher dose (8 g/day vs 1g/day). It is administered at an earlier and more vulnerable stage of pregnancy. Our study used physiological and supraphysiological concentrations of anti-CMV medications and specifically selected placental tissues from 10–13 weeks’ gestation to mirror the current recommendations on the prevention of first-trimester CMV infection in clinical care (4, 6, 30).

Clinical trials are currently evaluating the transplacental transfer of letermovir (NCT04732260) and its effectiveness in reducing fetal CMV infection (NCT05446571). While our work suggests possible alterations in Wnt and EMT gene expression, these do not appear to translate into impairment of placental viability or outgrowth. No studies currently address maribavir use in pregnant individuals, but our findings provide preliminary evidence that it may be safe for first-trimester placental function.

A key strength of this study is that it focuses on the first-trimester placenta. The risk of adverse consequences for the fetus is highest if the mother contracts primary CMV infection during the preconception period or the first trimester of pregnancy (31). Therefore, our study addresses the physiologically relevant window of the first trimester, rather than cell line models.

However, our research does not address the impact of first-trimester anti-CMV therapy on fetal teratogenesis or postnatal outcomes. Studies in animal models and pregnancy exposure registries are needed to address these gaps. Further research is currently underway to test the safety and efficacy of these agents in the same models in the presence of CMV infection.

## Conclusions

Our study provides the first data from first trimester primary human placental samples to support the safety of high-dose maternal valaciclovir treatment during the first trimester. Letermovir and maribavir, while not overtly detrimental to cell survival or villous outgrowth, may affect Wnt and EMT signalling pathways that are important for placental development. Future work should verify these findings in models of infected tissue and in larger clinical studies to guide the safe use of novel antivirals during early pregnancy.

## Notes

### Competing Interest Statement

The authors have declared no competing interest.

### Funding

- Northern Health
- University of Melbourne

## References

1. Ssentongo P, Hehnly C, Birungi P, Roach MA, Spady J, Fronterre C, et al. Congenital Cytomegalovirus Infection Burden and Epidemiologic Risk Factors in Countries With Universal Screening: A Systematic Review and Meta-analysis. JAMA Netw Open. 2021;4(8):e2120736.

2. Chatzakis C, Ville Y, Makrydimas G, Dinas K, Zavlanos A, Sotiriadis A. Timing of primary maternal cytomegalovirus infection and rates of vertical transmission and fetal consequences. Am J Obstet Gynecol. 2020;223(6):870-83.e11.

3. Zammarchi L, Tomasoni LR, Liuzzi G, Simonazzi G, Dionisi C, Mazzarelli LL, et al. Treatment with valacyclovir during pregnancy for prevention of congenital cytomegalovirus infection: a real-life multicenter Italian observational study. American Journal of Obstetrics & Gynecology MFM. 2023;5(10):101101.

4. Faure-Bardon V, Fourgeaud J, Stirnemann J, Leruez-Ville M, Ville Y. Secondary prevention of congenital cytomegalovirus infection with valacyclovir following maternal primary infection in early pregnancy. Ultrasound Obstet Gynecol. 2021;58(4):576–81.

5. Chatzakis C, Shahar-Nissan K, Faure-Bardon V, Picone O, Hadar E, Amir J, et al. The effect of valacyclovir on secondary prevention of congenital cytomegalovirus infection, following primary maternal infection acquired periconceptionally or in the first trimester of pregnancy. An individual patient data meta-analysis. Am J Obstet Gynecol. 2024;230(2):109-17.e2.

6. Shahar-Nissan K, Pardo J, Peled O, Krause I, Bilavsky E, Wiznitzer A, et al. Valaciclovir to prevent vertical transmission of cytomegalovirus after maternal primary infection during pregnancy: a randomised, double-blind, placebo-controlled trial. The Lancet. 2020;396(10253):779–85.

7. Leruez-Ville M, Chatzakis C, Lilleri D, Blazquez-Gamero D, Alarcon A, Bourgon N, et al. Consensus recommendation for prenatal, neonatal and postnatal management of congenital cytomegalovirus infection from the European congenital infection initiative (ECCI). The Lancet Regional Health – Europe. 2024;40.

8. Leruez-Ville M, Foulon I, Pass R, Ville Y. Cytomegalovirus infection during pregnancy: state of the science. Am J Obstet Gynecol. 2020;223(3):330–49.

9. Khorami-Sarvestani S, Vanaki N, Shojaeian S, Zarnani K, Stensballe A, Jeddi-Tehrani M, et al. Placenta: an old organ with new functions. Front Immunol. 2024;15:1385762.

10. Gude NM, Roberts CT, Kalionis B, King RG. Growth and function of the normal human placenta. Thrombosis Research. 2004;114(5):397–407.

11. Gupta SK, Malhotra SS, Malik A, Verma S, Chaudhary P. Cell Signaling Pathways Involved During Invasion and Syncytialization of Trophoblast Cells. American Journal of Reproductive Immunology. 2016;75(3):361–71.

12. Dietrich B, Haider S, Meinhardt G, Pollheimer J, Knöfler M. WNT and NOTCH signaling in human trophoblast development and differentiation. Cell Mol Life Sci. 2022;79(6):292.

13. Chronopoulou E, Koika V, Tsiveriotis K, Stefanidis K, Kalogeropoulos S, Georgopoulos N, et al. Wnt4, Wnt6 and β-catenin expression in human placental tissue - is there a link with first trimester miscarriage? Results from a pilot study. Reprod Biol Endocrinol. 2022;20(1):51.

14. Nayeem SB, Arfuso F, Dharmarajan A, Keelan JA. Role of Wnt signalling in early pregnancy. Reprod Fertil Dev. 2016;28(5):525–44.

15. Pollheimer J, Loregger T, Sonderegger S, Saleh L, Bauer S, Bilban M, et al. Activation of the canonical wingless/T-cell factor signaling pathway promotes invasive differentiation of human trophoblast. Am J Pathol. 2006;168(4):1134–47.

16. Sonderegger S, Husslein H, Leisser C, Knöfler M. Complex expression pattern of Wnt ligands and frizzled receptors in human placenta and its trophoblast subtypes. Placenta. 2007;28 Suppl A(Suppl A):S97–102.

17. Guo R, Xing QS. Roles of Wnt Signaling Pathway and ROR2 Receptor in Embryonic Development: An Update Review Article. Epigenet Insights. 2022;15:25168657211064232.

18. Vićovac L, Aplin JD. Epithelial-mesenchymal transition during trophoblast differentiation. Acta Anat (Basel). 1996;156(3):202–16.

19. Nakaya Y, Sheng G. EMT in developmental morphogenesis. Cancer Lett. 2013;341(1):9–15.

20. J Ed, Pollheimer J, Yong HE, Kokkinos MI, Kalionis B, Knöfler M, et al. Epithelial-mesenchymal transition during extravillous trophoblast differentiation. Cell Adh Migr. 2016;10(3):310–21.

21. Kokkinos MI, Murthi P, Wafai R, Thompson EW, Newgreen DF. Cadherins in the human placenta – epithelial–mesenchymal transition (EMT) and placental development. Placenta. 2010;31(9):747–55.

22. Du L, Kuang L, He F, Tang W, Sun W, Chen D. Mesenchymal-to-epithelial transition in the placental tissues of patients with preeclampsia. Hypertension Research. 2017;40(1):67–72.

23. Leruez-Ville M, Ghout I, Bussières L, Stirnemann J, Magny J-F, Couderc S, et al. In utero treatment of congenital cytomegalovirus infection with valacyclovir in a multicenter, open-label, phase II study. American Journal of Obstetrics and Gynecology. 2016;215(4):462.e1-.e10.

24. Faure Bardon V, Peytavin G, Lê MP, Guilleminot T, Elefant E, Stirnemann J, et al. Placental transfer of Letermovir & Maribavir in the ex vivo human cotyledon perfusion model. New perspectives for in utero treatment of congenital cytomegalovirus infection. PLoS One. 2020;15(4):e0232140.

25. Hamilton ST, Marschall M, Rawlinson WD. Investigational Antiviral Therapy Models for the Prevention and Treatment of Congenital Cytomegalovirus Infection during Pregnancy. Antimicrob Agents Chemother. 2020;65(1).

26. de Alwis N, Beard S, Binder NK, Pritchard N, Kaitu’u-Lino TuJ, Walker SP, et al. Placental OLAH Levels Are Altered in Fetal Growth Restriction, Preeclampsia and Models of Placental Dysfunction. Antioxidants. 2022;11(9):1677.

27. Kaitu’u-Lino TJ, Tong S, Beard S, Hastie R, Tuohey L, Brownfoot F, et al. Characterization of protocols for primary trophoblast purification, optimized for functional investigation of sFlt-1 and soluble endoglin. Pregnancy Hypertens. 2014;4(4):287–95.

28. MacDougall C, Guglielmo BJ. Pharmacokinetics of valaciclovir. Journal of Antimicrobial Chemotherapy. 2004;53(6):899–901.

29. Preibisch S, Saalfeld S, Tomancak P. Globally optimal stitching of tiled 3D microscopic image acquisitions. Bioinformatics. 2009;25(11):1463–5.

30. Pasternak B, Hviid A. Use of Acyclovir, Valacyclovir, and Famciclovir in the First Trimester of Pregnancy and the Risk of Birth Defects. JAMA. 2010;304(8):859–66.

31. Rybak-Krzyszkowska M, Górecka J, Huras H, Massalska-Wolska M, Staśkiewicz M, Gach A, et al. Cytomegalovirus Infection in Pregnancy Prevention and Treatment Options: A Systematic Review and Meta-Analysis. Viruses. 2023;15(11).

